# DNA methylation regulates transcriptional homeostasis of algal endosymbiosis in the coral model Aiptasia

**DOI:** 10.1101/213066

**Authors:** Yong Li, Yi Jin Liew, Guoxin Cui, Maha J. Cziesielski, Noura Zahran, Craig T. Michell, Christian R. Voolstra, Manuel Aranda

## Abstract

The symbiotic relationship between cnidarians and dinoflagellates is the cornerstone of coral reef ecosystems. Although research is focusing on the molecular mechanisms underlying this symbiosis, the role of epigenetic mechanisms, which have been implicated in transcriptional regulation and acclimation to environmental change, is unknown. To assess the role of DNA methylation in the cnidarian-dinoflagellate symbiosis, we analyzed genome-wide CpG methylation, histone associations, and transcriptomic states of symbiotic and aposymbiotic anemones in the model system Aiptasia. We find methylated genes are marked by histone H3K36me3 and show significant reduction of spurious transcription and transcriptional noise, revealing a role of DNA methylation in the maintenance of transcriptional homeostasis. Changes in DNA methylation and expression show enrichment for symbiosis-related processes such as immunity, apoptosis, phagocytosis recognition and phagosome formation, and unveil intricate interactions between the underlying pathways. Our results demonstrate that DNA methylation provides an epigenetic mechanism of transcriptional homeostasis during symbiosis.

## Introduction

Coral reefs are ecologically important marine ecosystems, which cover less than 0.2% of our oceans but sustain an estimated ~25% of the world’s marine species and 32 of 33 animal phyla (Spalding and Grenfell 1997; Davidson 2002; Sylvain 2006). Coral reefs are also economically important by providing food and livelihood opportunities to at least 500 million people; worldwide, they have a net present value of almost USD 800 billion, and they generate USD 30 billion in net economic benefits annually (Sylvain 2006). Unfortunately, these ecosystems are under severe threat from anthropogenic stressors including global warming and water pollution, among others, which can cause coral bleaching (loss of intracellular endosymbionts from coral) and overall coral reef decline. Despite increasing efforts on studying the mechanisms underlying the regulation and environmental stress related breakdown of this symbiotic association (Davy et al. 2012; Meyer and Weis 2012), we still lack knowledge on basic molecular processes, for instance whether epigenetic mechanisms are involved in symbiosis regulation and could potentially contribute to increased resilience in response to environmental stress as reported in other organisms (Rando and Verstrepen 2007; Lämke and Bäurle 2017).

DNA methylation plays an important role in many biological processes of plants and animals (Bird 2002; Suzuki and Bird 2008; He et al. 2011; Jones 2012). It has been proposed as a mechanism for organisms to adjust their phenotype in response to their environment in order to optimize organismal response to changing environmental conditions (Richards 2006; Rando and Verstrepen 2007). For instance, recent findings in mice show an important function for DNA methylation in inhibiting spurious transcription along the gene body, allowing for reduction of nonsense transcripts from highly expressed loci (Neri et al. 2017). Similar functions have also been proposed in plants, suggesting a general role of DNA methylation in the maintenance of transcriptional homeostasis (Zilberman 2017).

Several studies on DNA methylation in cnidarians have been published recently (Dixon et al. 2016; Putnam et al. 2016), however, its role and function in cnidarians is, at present, mainly unknown (Torda et al. 2017). The sea anemone Aiptasia is an emerging model to study the cnidarian-dinoflagellate symbiosis. Like corals, it establishes a stable but temperature sensitive symbiosis with dinoflagellates of the genus *Symbiodinium* but, unlike corals, can also be naturally maintained in an aposymbiotic state. This, compounded with its ease of culture, provides a tractable system to study the molecular mechanism underlying symbiosis without the impeding stress responses associated with coral bleaching stress (Voolstra 2013; Baumgarten et al. 2015).

Using the model system Aiptasia (strain CC7, sensu *Exaiptasia pallida*), we obtained whole-genome CpG DNA methylation, ChIP-Seq and RNA-Seq data from aposymbiotic (Apo) and symbiotic (Sym) individuals to study the function of DNA methylation in transcriptional regulation and its role in the cnidarian-dinoflagellate symbiosis.

## Results

### Aiptasia DNA Methylation patterns change with symbiotic states

To assess changes in DNA methylation in response to symbiosis, we performed whole-genome bisulfite sequencing with an average coverage of 53× per individual on 12 anemones, providing 6 biological replicates per treatment (symbiotic vs. aposymbiotic). Methylation calling using the combined dataset identified 710,768 CpGs (6.37% of all CpGs in Aiptasia genome), i.e. methylated sites in the Aiptasia genome. Notably, the percentage of CpGs is much lower than in mammals (60–90%) (Tucker 2001), but comparable to the coral *Stylophora pistillata* (7%) (Liew et al. 2017). We identified 10,822 genes (37% of all 29,269 gene models identified in the Aiptasia genome) with at least 5 methylated positions that were subsequently defined as methylated genes. On average, these genes had 18.4% CpGs methylated, 3-fold higher than the average methylation density across the entire genome (Chi-squared test *p* value < 2.2 × 10^−16^) and 167-fold higher than the methylation levels in non-coding regions. These findings indicate that the distribution of CpG methylation is non-random and mainly located in gene bodies, similar to corals (Dixon et al. 2017; Liew et al. 2017) and other invertebrate species (Feng et al. 2010; Gavery and Roberts 2010; Wang et al. 2013; Gonzalez-Romero et al. 2017).

To analyze the relationship between methylation density (percentage of CpGs) and gene density (the number of genes per 10,000 bp), we ran a sliding window (window size: 40 kb, step: 30 kb) and visualized the results in a Circos plot (Fig. S1) (Krzywinski et al. 2009). The correlation of CpG content and distribution of methylation showed a negative correlation (Pearson correlation coefficient: *r* = −0.31, *p* value < 2.2 × 10^−16^) suggesting that methylation tends to preferentially occur in CpG-poor regions (Fig. S2). Gene density had a positive correlation with methylation density (*r* = 0.21, *p* value < 2.2 × 10^−16^) consistent with the finding that methylation is predominantly located in gene bodies (Fig. 1). We also observed that within gene bodies, introns showed significantly higher methylation densities than exons (Fig. 1B).

**Fig. 1.**
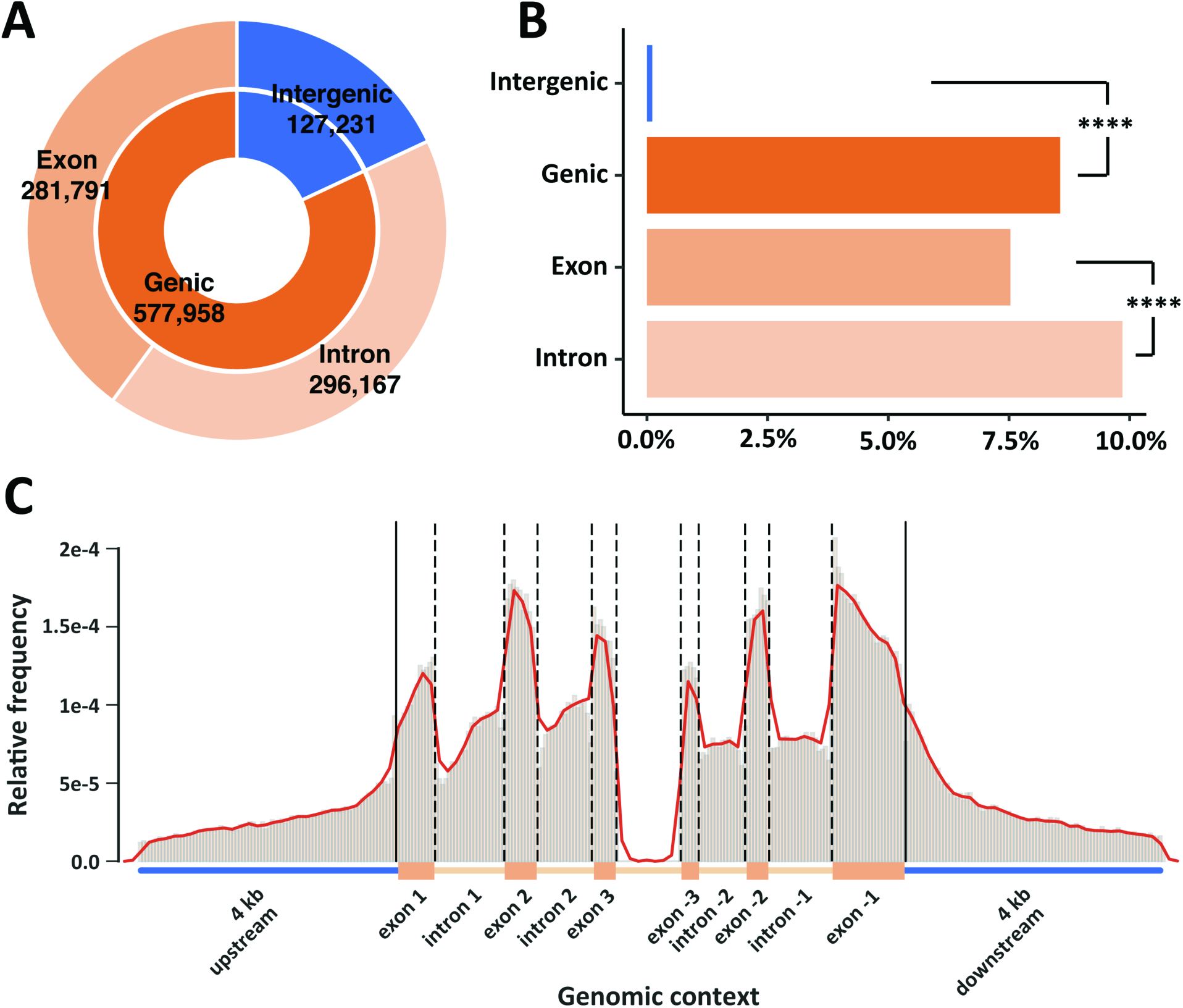
DNA methylation landscape. (**A**) Distribution of methylated CpG across intergenic (18%), genic (82%), intronic (42%) and exonic (40%) regions in the Aiptasia genome. (**B**) Normalized percentage of methylated CpGs in different regions. Chi-squared test shows significant differences between intergenic and genic regions, and between exons and intron (****p<0.0001). (**C**) Relative frequencies of methylated positions across a normalized gene model.

### Methylated genes are marked by H3K36me3

Analysis of methylation patterns (see above) within gene bodies showed rapidly increasing methylation levels after the transcription start site (TSS) that are maintained before slowly decreasing towards the transcription termination site (TTS) (Fig. S3A). Interestingly, we found that gene body methylation in Aiptasia is positively correlated with expression (Fig. 2A), suggesting that DNA methylation either increases the expression of genes or that DNA methylation is increased as a consequence of transcription whereby increased expression results in methylation of the respective loci. The latter interpretation would be in line with recent findings in mouse embryonic stem cells (Neri et al. 2017), which demonstrated that gene body methylation is established and maintained as a result of active transcription by RNA polymerase II (Pol II) and recruitment of the histone modifying protein SetD2 that trimethylates histone H3 at lysine 36 (H3K36me3). This histone mark is specifically bound via the PWWP domain present in the DNA methyltransferase Dnmt3b, which in turn methylates the surrounding DNA accordingly, resulting in the inhibition of transcription initiation from cryptic promoters within the gene body and thus a significant reduction of spurious transcription.

**Fig. 2.**
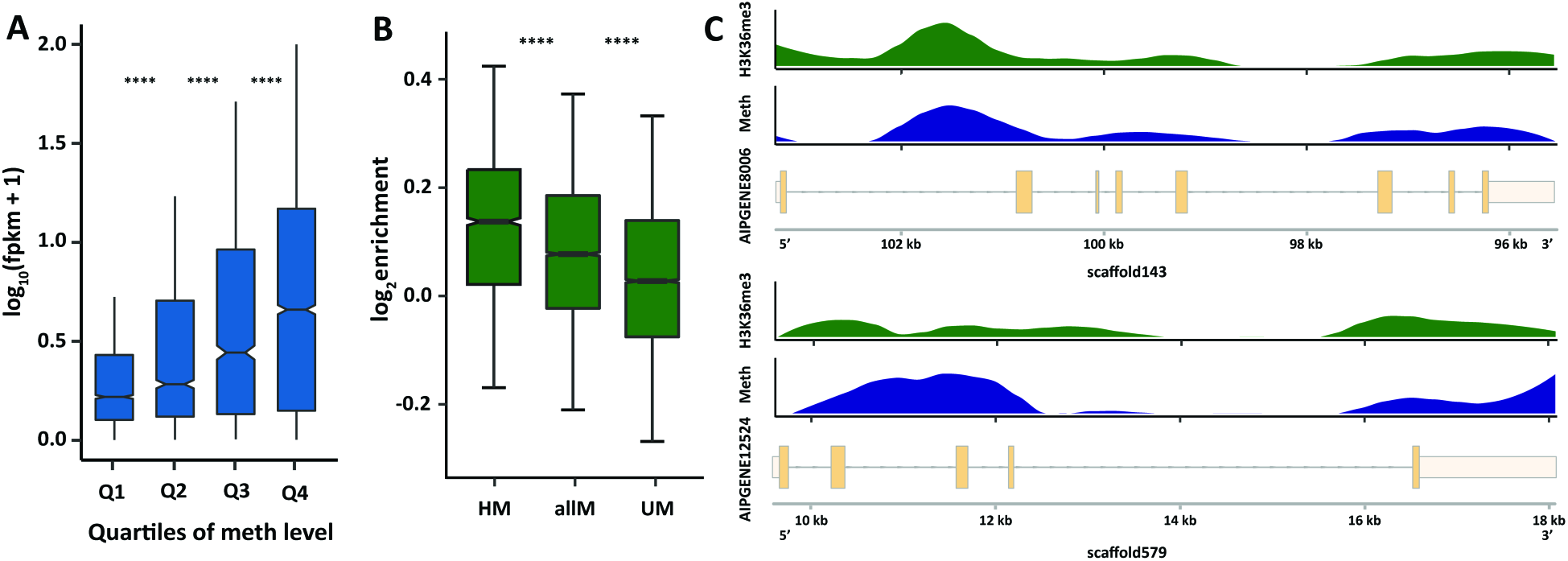
DNA methylation is associated with higher expression. (**A**) Gene expression is positively correlated with median methylation levels, *t*-test *p* values are 7.65 × 10^−21^, 3.75 × 10^−14^ and 1.75 × 10^−13^ for the first quartile (Q1) and the second quartile (Q2) of methylation levels, Q2 and Q3, and Q3 and Q4, respectively. (**B**) ChIP-Seq analysis of H3K36me3 signals show significant enrichment in methylated genes (*t*-test *p* values: 2.48 × 10^−20^ for highly methylated genes (HM) and all methylated genes (allM), and 1.06 × 10^−72^ for unmethylated genes (UM) and allM). Highly methylated genes show the strongest enrichment with H3K36me3 followed by all methylated genes. In contrast unmethylated genes show only weak enrichment of H3K36me3 over input controls. (**C**) Distribution of H3K36me3 enrichment and DNA methylation levels across two exemplary gene models. H3K36me3 and DNA methylation show coinciding distribution patterns over genes.

Analysis of the Aiptasia gene set identified a DNMT3 gene (AIPGENE24404) that also encodes a PWWP domain as reported for the mouse homolog. In order to test if the mechanism previously described in mice is conserved in Aiptasia, we performed a ChIP-Seq experiment using a validated antibody against H3K36me3 (Fig. S4-S5). As predicted, our analysis confirmed a significantly higher association of H3K36me3 with methylated genes (*p* = 2.48 × 10^−20^ for highly methylated genes and all methylated genes, Fig. 2B and C) suggesting that DNA methylation in Aiptasia might indeed be a consequence of expression. We then analyzed if methylated genes also exhibited significantly lower levels of spurious transcription in Aiptasia. Analysis of transcriptional profiles of methylated and unmethylated genes indeed showed significantly lower levels of spurious transcription along the gene body of methylated genes (*p* < 2 × 10^−6^, Fig. 3A).

**Fig. 3.**
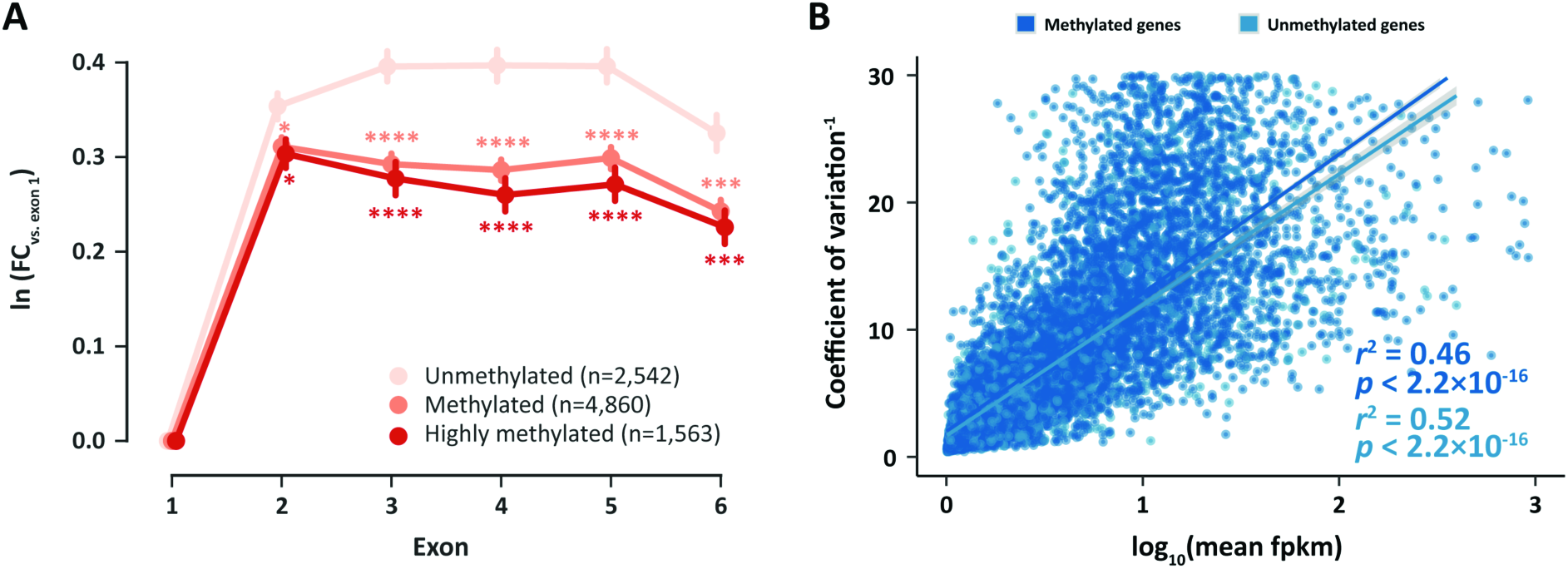
DNA methylation regulates transcriptional homeostasis. (A) Spurious transcription in gene bodies is significantly lower in methylated and highly methylated genes. The *y*-axis shows the natural logarithm of the coverage fold change of exons 1–6 vs. exon 1. *: *p* < 0.05; **: *p* < 0.01; ***: *p* < 0.001; ****: *p* < 0.0001. (B) There is a linear relationship between the inverse of transcriptional noise (CV^−1^) and log expression level (log_10_fpkm). Given same expression level, methylated genes always show lower levels of transcriptional noise. For methylated genes, n = 8,561, *r*^2^ = 0.46, *p* < 2.2 × 10^−16^, for unmethylated genes, n = 2,491, *r*^2^ = 0.52, *p* < 2.2 × 10^−16^.

A dampening effect of DNA methylation on transcription was also observed with regard to transcriptional noise similar to findings in the coral *Stylophora pistillata* (Liew et al. 2017). Regression analysis of median methylation levels and the coefficient of transcriptional variation of genes showed that, given the same expression level, methylated genes always exhibited lower levels of transcriptional variation (Fig. 3B).

### DNA methylation regulates transcriptional homeostasis during symbiosis

Based on our previous findings, we investigated if DNA methylation might also be involved in the regulation of symbiosis by identifying differentially methylated genes (DMGs) between symbiotic and aposymbiotic Aiptasia. Comparison of DNA methylation patterns using Principal Component Analysis (PCA) clearly separated symbiotic and aposymbiotic individuals by the first principal component, which accounted for ~18% of the variance (Fig. 4 and Fig. S6). This analysis echoed the findings from a PCA analysis on gene expression where symbiosis state was separated by the second principal component accounting for ~25% of the variance (Fig. 4B) (Venables and Ripley 2002) and highlighted that specific changes in DNA methylation patterns occurred in response to symbiosis. Analysis of DNA methylation and expression profiles using correlation analyses further confirmed this finding, providing additional evidence that the observed changes were indeed treatment specific (Fig. S6).

**Fig. 4.**
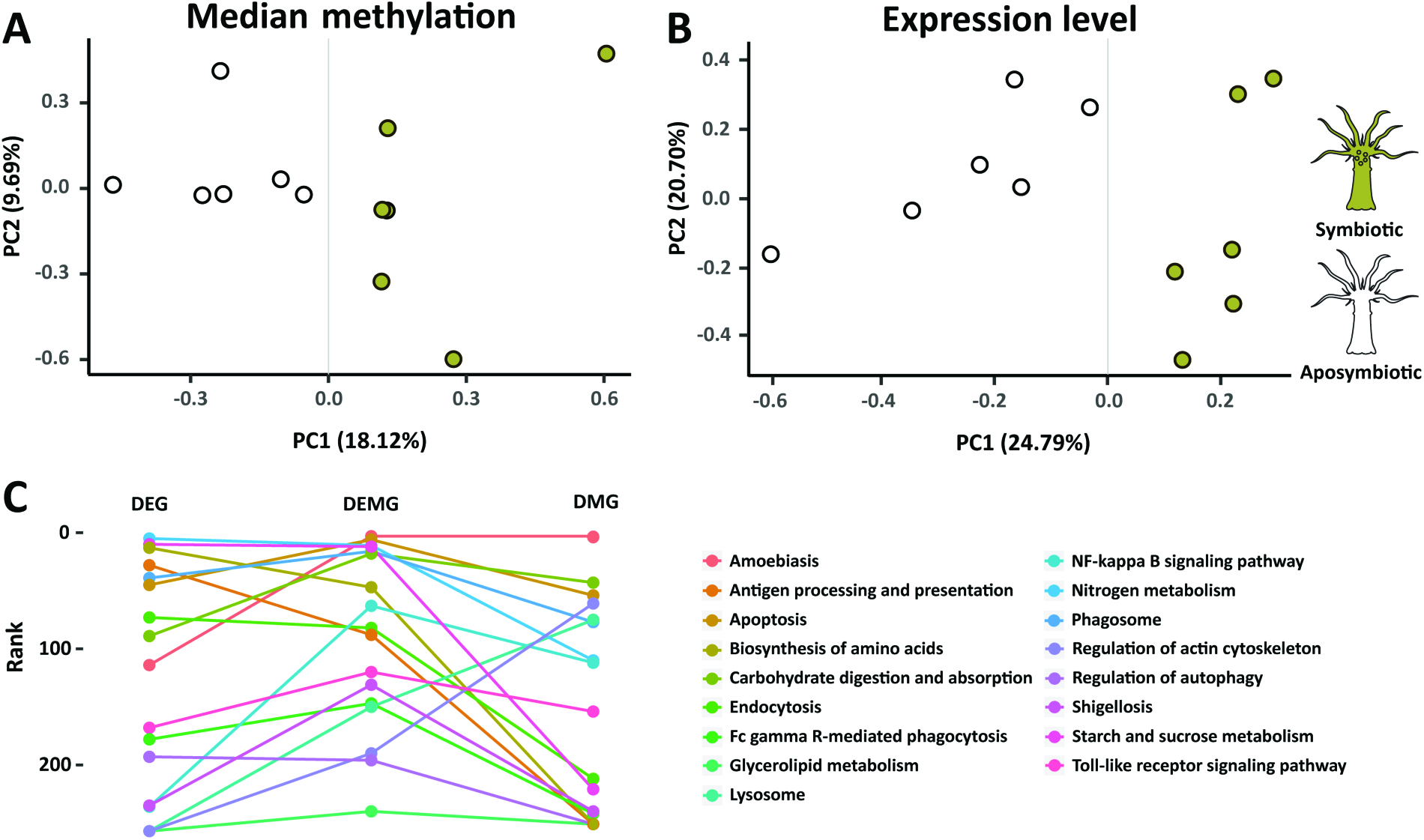
PCA and KEGG pathway enrichment analysis. (**A**, **B**) PCA (Principal Component Analysis) of gene expression and median methylation levels of Aiptasia genes. Both gene expression and DNA methylation separate samples by symbiosis state. (**C**) KEGG pathway enrichment analysis. The combined sets of differentially expressed and differentially methylated genes (DEMG) provides significant lower *p* values (front ranks) for symbiosis related pathways.

Subsequently we analyzed changes in DNA methylation and gene expression between symbiotic and aposymbiotic Aiptasia to assess their correlation on potential biological functions in symbiosis. We determined differentially methylated genes using a generalized linear model from Foret *et al.* (Foret et al. 2012) that was modified to allow for replicate-aware analysis. This approach identified 2,133 DMGs (FDR ≤ 0.05, Supplement Table S1) that specifically changed their methylation status in response to symbiosis. To verify these results, we sequenced a subset of 14 DMGs using bisulfite PCRs. The results show a strong correlation (*r*^2^ = 0.815 and *p* = 1×10^−5^ for Apo, *r*^2^ = 0.922 and *p* = 5.2 × 10^−8^ for Sym) to our WGBS and confirm the observed methylation changes within these loci (Fig. S7).

Analysis of gene expression changes in the same 12 samples (i.e., 6 symbiotic and 6 aposymbiotic anemones) identified 1,278 differentially expressed genes (DEGs, FDR ≤ 0.05, Supplement Table S2), of which 14 genes were subsequently confirmed via qPCR (Fig. S8). However, analysis of the overlap between DMGs and DEGs showed only 103 genes that were shared, suggesting that differentially expressed genes are not necessarily the same cohort of genes that are differentially methylated. Functional enrichment analyses based on Gene Ontology (GO) and Kyoto Encyclopedia of Genes and Genomes (KEGG) pathways of all DMGs and DEGs identified several symbiosis relevant functions and pathways in both groups (Supplement Table S3-S10).

Based on the finding that gene body DNA methylation is likely a consequence of active transcription, we hypothesized that changes in DNA methylation patterns might also provide a record of transcriptional activity over longer periods of time. We therefore tested if differential methylation and acute transcriptional changes, obtained from our RNA-Seq analysis, provide a complementary view of the processes underlying symbiosis. For this we compared enrichment of symbiosis-specific pathways across the sets of 2,133 DMGs, 1,278 DEGs, and the combined set of both DMGs and DEGs (3,308 DEMGs). Interestingly, we observed that the combined data set (DEMGs) provided significantly lower p-values for previously identified symbiosis-related pathways, including apoptosis, phagosome formation, nitrogen metabolism, and arginine biosynthesis, among others (paired *t*-test: DEMG vs. DEG *p* = 0.015; DEMG vs. DMG *p* = 0.009) (Fig. 4C and Supplement Table S11). This suggested that changes in methylation and transcription indeed provide complementary information with regard to transcriptional adjustments in response to symbiosis.

### DMGs and DEGs are involved in all stages of symbiosis

Analysis of the combined DMG and DEG gene set showed significant enrichment of genes involved in the distinct phases of symbiosis, that is symbiosis establishment, maintenance, and breakdown (Davy et al. 2012). Using an integrated pathway analysis based on known molecular interactions between proteins we found that these processes are linked through several DMGs and/or DEGs (Fig. S9 and Fig. S10, and see Supplement Table S11-S12 and Supplementary discussion).

For instance, we found numerous symbiosis-related receptors to respond to symbiosis on a transcriptional and/or methylation level (Fig. S9), including C-type lectins (Fig. S9.3), Toll-like receptors (Fig. S9.5), and the scavenger receptor SRB1 (Fig. S9.2) that has previously been implicated in symbiont recognition in the sea anemone *Anthopleura elegantissima* (Rodriguez-Lanetty et al. 2006; Neubauer et al. 2016). Following symbiont recognition, we also found several known engulfment and sorting-related genes to change in methylation and/or expression such as Rab5 (Fig. S9.10), sorting nexin (Fig. S9.17), Rac1 (Fig. S9.6), the lysosomal-associated membrane protein 1/2 (Fig. S9.22), and many genes related to the cytoskeleton and movement (Fig. S9.33-39).

As expected in a metabolic symbiosis (Muscatine 1990; Davy et al. 2012) we also identified a large number of genes involved in nutrient exchange. These included genes involved in the provision of inorganic carbon in the form of CO_2_ or bicarbonate (HCO_3_^-^) to fuel symbiont driven photosynthesis (Rädecker et al. 2017) (Fig. S10.1) as well as genes involved in the exchange of fixed carbon in the form of lipids (Fig. S10.11), sugars and amino acids (Fig. S10.10, S10.4) (Oakley et al. 2016). Concordantly, we also found that genes involved in nitrogen acquisition, such as ammonium transporter (Fig. S10.2) and genes involved in glutamate metabolism (Fig. S10.5-7), respond to symbiosis.

Finally, our analysis also highlighted genes putatively involved in the expulsion or degradation of symbionts in response to environmental stress or as a means to control symbiont densities. Autophagy is of interest in this regard because it links to other membrane trafficking pathways and to apoptosis, and evidence suggests that autophagy also plays a role in removal of symbionts during bleaching (Dunn et al. 2007; Downs et al. 2009). Intracellular degradation of the symbiont is a result of reengagement of the phagosomal maturation process or autophagic digestion of the symbiont by the host cell (Davy et al. 2012), and we find both apoptosis-and autophagy-related genes to significantly change in their methylation and/or expression level. These include the apoptosis genes RAC serine/threonine-protein kinase (Fig. S9.25), Caspase 7 (Fig. S9.31), Caspase 8 (CASP8) (Fig. S9.30), Nitric oxide synthase (Fig. S9.21) and Bcl2 (Fig. S9.27), as well as the Autophagy proteins 5 and 10 (Fig. S9.14-15), among others.

## Discussion

To assess the role of CpG methylation in the cnidarian-dinoflagellate symbiosis, we undertook a global analysis of changes in the DNA methylomes and transcriptomes of aposymbiotic and symbiotic Aiptasia. In contrast to their vertebrate counterparts, only 6.37% of the CpGs in the Aiptasia genome are methylated, but their distribution is highly non-random (*p* < 3 × 10^−300^) and that methylated CpGs are most highly localized in gene bodies (18.4% of CpGs). Analysis of the distribution of the histone modification H3K36me3 further showed significant enrichment of this epigenetic mark in methylated genes, echoing findings in mammals and invertebrates (Nanty et al. 2011). More importantly, we find that methylated genes show significant reduction of spurious transcription and transcriptional noise (Fig. 2B), suggesting that both the underlying mechanism of epigenetic crosstalk as well as the biological function of DNA methylation is evolutionarily conserved throughout metazoans. These results highlight a tight interaction of transcription and epigenetic mechanisms in optimizing gene expression in response to changing transcriptional needs (Neri et al. 2017). Further support for such a role is provided by the analysis of differentially methylated and differentially expressed genes, which, when combined, showed significant increase in enrichment of symbiosis relevant processes. This suggests that DNA methylation and transcriptome analyses provide complementary views of cellular responses to symbiosis whereby methylation changes provide a transcriptional record of longer-term transcriptional adjustments.

While our analysis identified several genes, processes, and pathways previously reported to be involved in symbiosis, it further highlights their intricate molecular interactions. Symbiosis recognition, sorting and breakdown are interconnected processes, which is reflected in the observed changes in methylation and expression. The molecular machinery involved in phagosome maturation is tightly linked to autophagy and apoptosis enabling the host to respond to potential pathogen invasion but also to degrade and remove dead or unsuitable symbionts. This is strongly supported by immunofluorescence examinations of *Aiptasia pulchella* gastrodermal cell macerates, showing that Rab5 appears around healthy, newly ingested and already established *Symbiodinium*, but is replaced by Rab7 in heat-killed or DCMU-treated newly ingested *Symbiodinium*. Conversely, Rab7 is absent from untreated newly infected or already-established *Symbiodinium* (Chen et al. 2003; Chen et al. 2004).

Rab5 is also required for the exosomal release of CD63 (Baietti et al. 2012), which mediates the endocytotic sorting process and transport to lysosomes (Latysheva et al. 2006). This process is further regulated by Rac1 (Anitei et al. 2010) in conjunction with sorting nexin and the GTPase Rho, all of which were also identified in our analyses. The sorting of phagocytosed *Symbiodinium* is critical to symbiosis establishment as *Symbiodinium* cells are phagocytosed at the apical end and transported to the base of the cell, where they are protected from digestion. In contrast, *Symbiodinium* staying at the apical end of the cell are degraded (McAuley and Smith 1982).

Similar to the processes of symbiosis initiation and breakdown, we also found significant enrichment of genes involved in nutrient exchange and many of these transporters have previously been implicated in symbiosis maintenance (Davy et al. 2012; Lin et al. 2015). Notably, this also included genes involved in the transport and assimilation of ammonium. Nitrogen is a main limiting nutrient in coral reefs (Cook et al. 1992; Grover et al. 2008; Radecker et al. 2015), and the coral-dinoflagellates symbiosis has been proposed to increase the efficiency of nitrogen utilization by both partners (Wang and Douglas 1998b) whereby the underlying nature of this mechanism is currently debated (Wang and Douglas 1998a; Aranda et al. 2016).

## Conclusions

This study provides the first analysis of the function and role of DNA methylation in a symbiotic anthozoan. Our results show that the epigenetic crosstalk between the histone mark H3K36me3 and gene body methylation is conserved in cnidarians and reveal a role of gene body methylation in reducing of spurious transcription and transcriptional noise. Furthermore, we show that changes in DNA methylation patterns are specific to symbiosis and imply a functional in the establishment, maintenance, and breakdown of this important symbiotic association. Our findings therefore provide evidence for a role of DNA methylation as an epigenetic mechanism involved in the maintenance of transcriptional homeostasis during the cnidarian-dinoflagellate symbiosis. The premise that epigenetic mechanisms play a role in organismal acclimation warrants future experiments targeted to investigate if DNA methylation could also contribute to resilience through the epigenetic adjustment of transcription in response to environmental stress in Aiptasia and corals.

## Data availability

Sequencing data of Bis-Seq, RNA-Seq and ChIP-Seq were deposited in NCBI Sequence Read Archive (SRA) under BioProject codes PRJNA415358.

## Acknowledgements

Research reported in this publication was supported by funding from King Abdullah University of Science and Technology (KAUST). We thank Prof. John Pringle for the provision of the initial Aiptasia CC7 and *Symbiodinium* SSB01 cultures. We thank Sebastian Baumgarten for his help with establishing the Aiptasia cultures and critical reading of the manuscript.

## Author contributions

M.A. conceived and coordinated the project. Y.L., G.C., M.J.C. and N.Z. performed experiments. M.A., C.R.V. and Y.J.L. provided tools and/or data. C.T.M. constructed libraries for whole genome bisulfite sequencing, ChIP-Seq and RNA-Seq. Y.L., Y.J.L. and M.A. analyzed expression, methylation and ChIP-Seq data. M.A. and Y.L. wrote the manuscript with input from Y.J.L. and C.R.V. All authors read and approved the final manuscript.

## Supplementary Materials

### Materials and methods

#### *Exaiptasia pallida* Culture and DNA/RNA Extraction

*Exaiptasia pallida* of the clonal strain CC7 (Sunagawa et al. 2009) was used for this study. Anemones were maintained in polycarbonate tubs with autoclaved seawater at ~25 °C on a 12 h: 12 h light: dark cycle at 20-40 μmol m^−2^ s^−1^ light intensity and fed freshly hatched *Artemia nauplii* (brine-shrimp) approximately twice per week. To generate aposymbiotic anemones, animals were subjected to multiple cycles of cold-shock treatment and the photosynthesis inhibitor diuron (Sigma-Aldrich, St. Louis, MO) as described in Baumgarten (2015) (Baumgarten et al. 2015). Aposymbiotic anemones were kept individually in 15 ml autoclaved seawater in 6-well plates and inspected by fluorescence stereomicroscopy to confirm complete absence of dinoflagellates. In order to exclude potential batch effects as source of DNA methylation changes we first generated four separate batches of aposymbiotic anemones and maintained them for a period of 1 year before beginning of the experiment described below.

To generate symbiotic anemones, we then separately infected aposymbiotic CC7 individuals from each of the four aposymbiotic cultures described above using the compatible Clade B *Symbiodinium* strain SSB01, originally isolated from Aiptasia strain H2 (Xiang et al. 2013; Baumgarten et al. 2015). The four batches of symbiotic anemones were maintained for further 12 months under regular culture condition as described above. The corresponding four aposymbiotic cultures were maintained in darkness until 3 months before collection. For the last 3 months individuals from these aposymbiotic cultures were subjected to the same culture conditions as the symbiotic cultures in order to monitor for unwanted spontaneous re-establishment of symbiosis under light.

After the 12-month experimental period, we collected six biological replicates from each of the four aposymbiotic and symbiotic cultures (one additional replicate was taken from batches 1 and 2 of each treatment) for subsequent DNA and RNA extraction as described below.

For each treatment, 6 biological replicates, weighing 20-28 mg (wet weight), were extracted using the AllPrep DNA/RNA/miRNA Universal Kit (Qiagen, Hilden, Germany). The manufacturer’s protocol was followed with the omission of the optional step 4 (temporal storage at 4°C if not performing DNA purification immediately). DNA concentrations were determined using a Qbit dsDNA HS Assay Kit (Thermo Fisher Scientific, Waltham, MA). RNA concentrations and integrity were determined using a Bioanalyzer Nano RNA Kit (Agilent Technologies, Santa Clara, CA).

#### RNA-Seq and Bisulfite Sequencing

Directional mRNA libraries were produced using the NEBNext^®^ Ultra™ Directional RNA Library Prep Kit for Illumina^®^ (NEB) following the manufacturer’s protocol.

Bisulfite DNA libraries were prepared following a modified version of the NEBNext^®^ Ultra™ II DNA Library Prep Kit for Illumina^®^ (NEB). Methylated TruSeq Illumina^®^ adapters (Illumina) were used during the adapter ligation step followed by bisulfite conversion with the EpiTect Bisulfite kit (QIAGEN), with the following cycling conditions (95°C – 5 min, 60°C – 25 min, 95°C – 5 min, 60°C – 85 min, 95°C – 5 min, 60°C – 175 min, and 3 cycles of 95°C – 5 min, 60°C – 180 min, hold at 20°C ≤ 5 hours) (reference is Illumina Bisulfite).

The final libraries were enriched with the KAPA HiFi HotStart Uracil+ ReadyMix (2X) (KAPA Biosystems) following the standard protocol for bisulfite-converted NGS library amplification. Final libraries were quality checked using the Bioanalyzer DNA 1K chip (Agilent), and quantified using Qubit 2.0 (Thermo Fisher Scientific), and then pooled in equimolar ratios and sequenced on the HiSeq2000.

#### Identification of methylated CpGs

Sequencing of the 12 libraries (2 conditions, 6 biological replicates each) resulted in 819 million read pairs from 8 lanes of the Illumina HiSeq2000 platform. Adapters were trimmed from the raw sequences using cutadapt v1.8 (Martin 2011). Subsequently, trimmed reads were mapped to the *Exaiptasia pallida* genome (Baumgarten et al. 2015) using Bowtie2 v2.2.3 (Langmead and Salzberg 2012), and methylation calls was performed using Bismark v0.13 (Krueger and Andrews 2011).

Three filters were used to reduce false positives. Firstly, for each position with *k* methylated reads mapping to it, the probability of it occurring through sequencing error (i.e. unmethylated position appearing as methylated) was modelled using a binomial distribution B(*n, p*), where *n* is the coverage (methylated + unmethylated reads) and *p* the probability of sequencing error (set to 0.01). We kept positions with *k* methylated reads if P(X ≥ *k*) < 0.05 (post-FDR correction). Secondly, retained methylated positions had to have ≥ 1 methylated read in all six biological replicates of at least one growth condition. Finally, median coverage of retained positions across all 12 samples had to be ≥ 10.

#### Assignment of genomic context to methylated cytosines

Based on the gene annotation of the *Exaiptasia pallida* genome (GFF3 file) (Baumgarten et al. 2015) and the positional coordinates of the methylated cytosines produced by Bismark, we annotated every methylated cytosine based on the genomic context, including whether the methylated position resides in a genic or intergenic region, and the distances to the 5’ and 3’ end of each genomic feature (gene/intergenic region/exon/intron).

#### CpG bias

Methylated cytosines are frequently spontaneously deaminated to uracil which can be subsequently converted to thymine after DNA repair. As a result of this process, methylated CpGs are expected to decrease in abundance over evolutionary time, and the ratio of observed to expected CpGs (CpG O/E) has previously been used to predict putatively methylated and unmethylated genes (Suzuki et al. 2007; Wang and Leung 2008). CpG O/E of *Exaiptasia pallida* protein coding genes were calculated according to J. Zeng *et al* (Zeng and Yi 2010).

#### Identification of differentially methylated genes

Using the methylation level of aposymbiotic genes as a control, generalized linear models (GLMs) (Hastie and Pregibon 1992) were implemented in R (R Core Team 2016) to identify genes that were differentially methylated in the symbiotic treatment. The general formula used was:
glm(methylated, non_methylated ~ treatment * position, family=“binomial”)
where “methylated, non_methylated” was a two-column response variable denoting the number of methylated and non-methylated reads at a particular position. For predictor variables, “position” denoted relative position of the methylated site in the gene, while “treatment” denoted symbiotic or aposymbiotic conditions. Data from individual replicates were entered separately to assign equal weightage to each replicate, as pooling results in a disproportionate skew towards the replicate with the highest coverage. Genes with < 5 methylated positions were filtered out to reduce type I errors; and genes with FDR ≤ 0.05 were considered as differentially methylated genes (DMGs).

#### Identification of differentially expressed genes

RNA-Seq generated 889 million raw read pairs from six lanes on the Illumina Hiseq2000 platform. Adaptors, primers and low quality bases were removed from the ends of raw reads using Trimmomatic v0.33 (ILLUMINACLIP:TruSeq2-PE.fa:4:25:9 LEADING:28 TRAILING:28 SLIDINGWINDOW:4:30 MINLEN:50). The resulting trimmed reads were mapped to the <u>*Exaiptasia pallida*</u> genome using HISAT v2.0.1 (Kim et al. 2015) and transcripts were assembled based on the <u>*Exaiptasia pallida*</u> gene models (GFF3 file) using StringTie v1.2.2 (Pertea et al. 2015). Trinity (align_and_estimate_abundance.pl – Bowtie2 v2.2.7/RSEM v1.2.22/edgeR v3.10.5) (Robinson et al. 2010; Grabherr et al. 2011; Li and Dewey 2011; Langmead and Salzberg 2012; Haas et al. 2013) was run against the transcripts using trimmed reads for expression abundance estimation, then differentially expressed genes (DEGs) were identified with FDR ≤ 0.05.

#### PCA and correlation matrices

Median methylation levels and log FPKM (base 2) of genes were shifted to be zero centered and analyzed by <u>Principal Component Analysis</u> (PCA) using the prcomp function in R.

Using the same data we calculated correlation matrices (Pearson correlation coefficient) and clustered samples with hclust implemented in R using complete linkage and euclidean distance.

#### Spurious transcription analysis

Trimmed reads were mapped to the <u>*Exaiptasia pallida*</u> genome using HISAT2 v2.1.0 and mapping coverage per position was extracted using BEDtools v2.17.0. Coverage per exon was calculated and normalized across all 6 replicates (assuming every replicate had 1 million coverage in total), then average coverage ratios of exon 2 to 6 versus exon 1 per gene were calculated to determine spurious transcription levels.

#### GO enrichment of DMGs and DEGs

GO (Gene Ontology) (Ashburner et al. 2000) annotation was based on the previously published *Exaiptasia pallida* genome (Baumgarten et al. 2015). Functional enrichment of DMGs and DEGs was carried out with topGO respectively (Adrian Alexa 2016) using default settings. GO terms with *p* ≤ 0.05 were considered significant, and the occurrence of at least ≥ 5 times in the background set was additionally required for DMGs. Multiple testing correction was not applied to the resulting *p*-values as the tests are considered to be non-independent (Adrian Alexa 2016).

#### KEGG enrichment of DMGs and DEGs

KEGG (Kyoto Encyclopedia of Genes and Genomes)(Kanehisa and Goto 2000; Kanehisa et al. 2016) orthology (KO) annotation was carried out by combining the KEGG annotations provided in the original *Exaiptasia pallida* genome publications and a separate set of annotations based on the KAAS (KEGG Automatic Annotation Server, http://www.genome.jp/tools/kaas/) (parameters: GHOSTZ, Eukaryotes, Bi-directional Best Hit) (Moriya et al. 2007). A KEGG pathway enrichment analysis of both DMGs and DEGs was carried out using Fisher’s exact test and pathways with *p* ≤ 0.05 were considered significant.

#### Validation of gene expression changes from RNA-Seq by qPCR

Three randomly picked RNA libraries per treatment were used for qPCR validation of RNA-Seq results. cDNA was synthesized using Invitrogen SuperScript III First-Strand Synthesis SuperMix kit. A total of 14 genes were validated for differential expression using qPCR (Supplement Table S13-S15). RPS7, RPL11 and NDH5 were used as internal reference standards (Lehnert et al. 2014). qPCR was carried out using Invitrogen Platinum SYBR Green qPCR SuperMix-UDG kit on Applied Biosystems 7900HT Fast Real-Time PCR System. All protocols were strictly followed.

#### Validation of methylation changes using bisulfite PCR

Three randomly picked DNA libraries per treatment were used for methylation validation. Bisulfite conversion was done using the EZ-96 DNA Methylation-Gold Kit (Zymo Research). 18 genes were used to design primers, 14 of 18 obtained effective amplifications (Supplement Table S16), then the fragments were enriched by PCR amplification using Promega PCR Master Mix. Sequencing indices were added to enriched fragments using Illumina Nextera XT Index Kit. Enriched fragments were sequenced on the Illumina MiSeq platform. All protocols were strictly followed. 1,870x data per replicate were obtained, methylated CpGs were identified using Bismark as described above. The correlations between whole genome bisulfite conversion and bisulfite PCR were calculated using generalized linear model.

#### Chromatin Immunoprecipitation – ChIP

We used the Zymo-Spin ChIP Kit to conduct histone bound chromatin extraction, with minor adjustments to manufacturer’s protocol. Briefly, three biological replicates, each consisting of two symbiotic anemones, were used for this experiment. Each anemone was first washed with PBST (phosphate-buffered saline with 0.1% triton). Anemones were then fixed in 1X PBS with 1% formaldehyde for 15 minutes. To stop cross-linking reactions glycine was added to the solution and left to rest for 10 more minutes. Following manufacturer’s protocol, we centrifuged and washed whole anemones. We prepared the Nuclei Prep Buffer according to protocol and crushed the two anemones of each replicate together using a douncer for homogenization. Samples were then sonicated for 15 cycles on ice (15 sec ON, 30 sec cooling) to ensure fragmentation to 200-500 bp. Thereafter the protocol was followed without further modifications.

Immunoprecipitation was achieved using a target specific antibody to histone 3 lysine 36 tri-methylation (H3K36me3) (ab9050, Abcam), which has been validated in many eukaryotic species, including mouse(Soboleva et al. 2017), *Arabidopsis thaliana*(Wollmann et al. 2017), yeast(Janke et al. 2017) and zebrafish(Vastenhouw et al. 2010; Wu et al. 2011) *et al.* Comparison of Aiptasia histone H3 to the respective homologs from several species for which this antibody has been previously validated showed high conservation of the N-terminal tail containing position H3K36 (Fig. S4) whereby 100% conservation to the zebrafish homolog was observed.

Corresponding input controls for each of the 3 replicates were generated as suggested by the manufacturer. DNA fragment quality and quantity were confirmed using High Sensitivity DNA Reagents (Agilent Technologies, California, United States) on a bioanalyzer, after which ChIP libraries were constructed using the NEBNext^®^ ChIP-Seq Library Prep Master Mix Set (#E7645, New England Biolabs, Massachusetts, United States).

Sequencing resulted in 10M read pairs per replicate. These read pairs were trimmed using Trimmomatic and mapped to the *Exaiptasia pallida* genome using HISAT2 as described above. H3K36me3 enrichments were calculated as log_2_(average signal/average input control) for all genes, unmethylated genes and highly methylated genes (methylation level > 70 and methylation density > 40). P-values were calculated using t-test.

#### Antibody affinity validation through Western blotting

Total protein was extracted from a snap-frozen anemone crushed in 10% TCA (Trichloroacetic acid). The homogenized sample was left to incubate overnight at −20 °C to allow proteins to precipitate. The solution was centrifuged at 20,000g at 4 °C for 20 minutes to collect suspended proteins. The pellet was then washed three times in 80% acetone and then spun down again as previously. The final pellet was then air-dried for 10-15 minutes to remove residual acetone. The final protein was suspended in urea lysis buffer (7 M urea, 2 M thiourea, pH 7.5) by vortexing for 2 hours.

Samples were then prepared for western blot by adding 4x sampling buffer (0.38 M Tris base, 8% SDS, 4mM EDTA, 40% glycerol, 4mg/ml bromphenol blue) to a final concentration of 1X. After a 2 minute incubation at 90°C samples were ready to be run on a gel at 10-12 mA. The gel was transferred to a PVDF nitrocellulose membrane, rinsed with TBS buffer (150mM NaCl, 25mM Tris pH7.4, 0.1% Triton X-100) and blocked for 30 min at RT in TBS containing 5% fat-free powder milk. The primary antibody was diluted in TBS/milk and incubated on an undulating orbital shaker overnight at 4 °C. After three washes in TBS for 10 minutes each, the membrane was again blocked in TBS/milk for 20 minutes at RT before proceeding with secondary antibody staining. The horseradish peroxidase-linked-antibody (Anti-Rabbit IgG HRP conjugate W4011 and Anti-Mouse IgG HRP conjugate W4031, Promega. Wisconsin, United States) was diluted in TBS/milk (1:10000) and incubated for 2 hours at RT. After final triplicate 10 minute washes in TBS, membranes were developed.

**Fig. S1.**
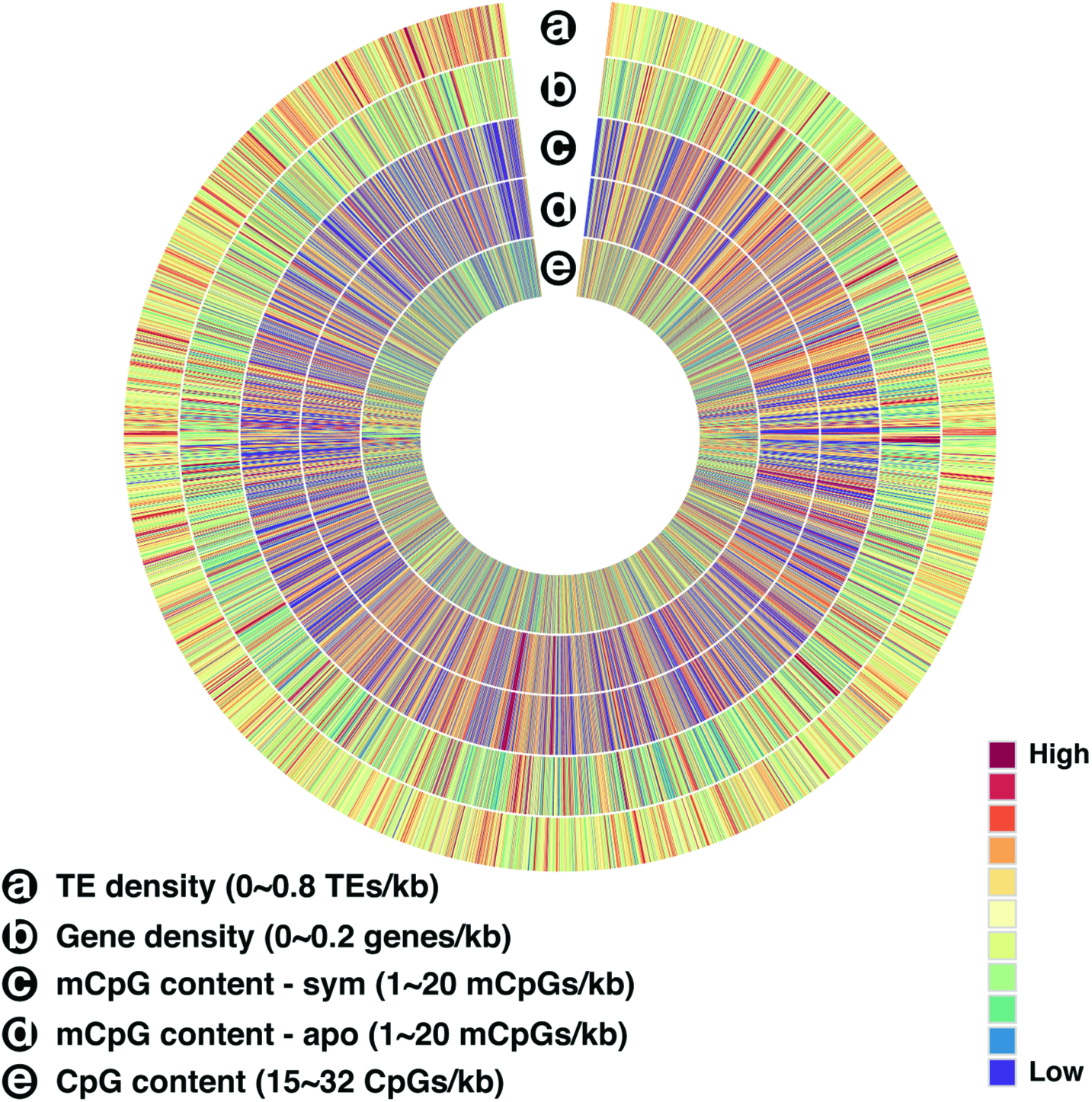
Circos visualization of different data at the genome-wide level. (**a**) TE density. (**b**) Gene density. (**c**) Fraction of methylated CpGs in symbiotic treatment. (**d**) Fraction of methylated CpGs in aposymbiotic treatment. (**e**) CpG content.

**Fig. S2.**
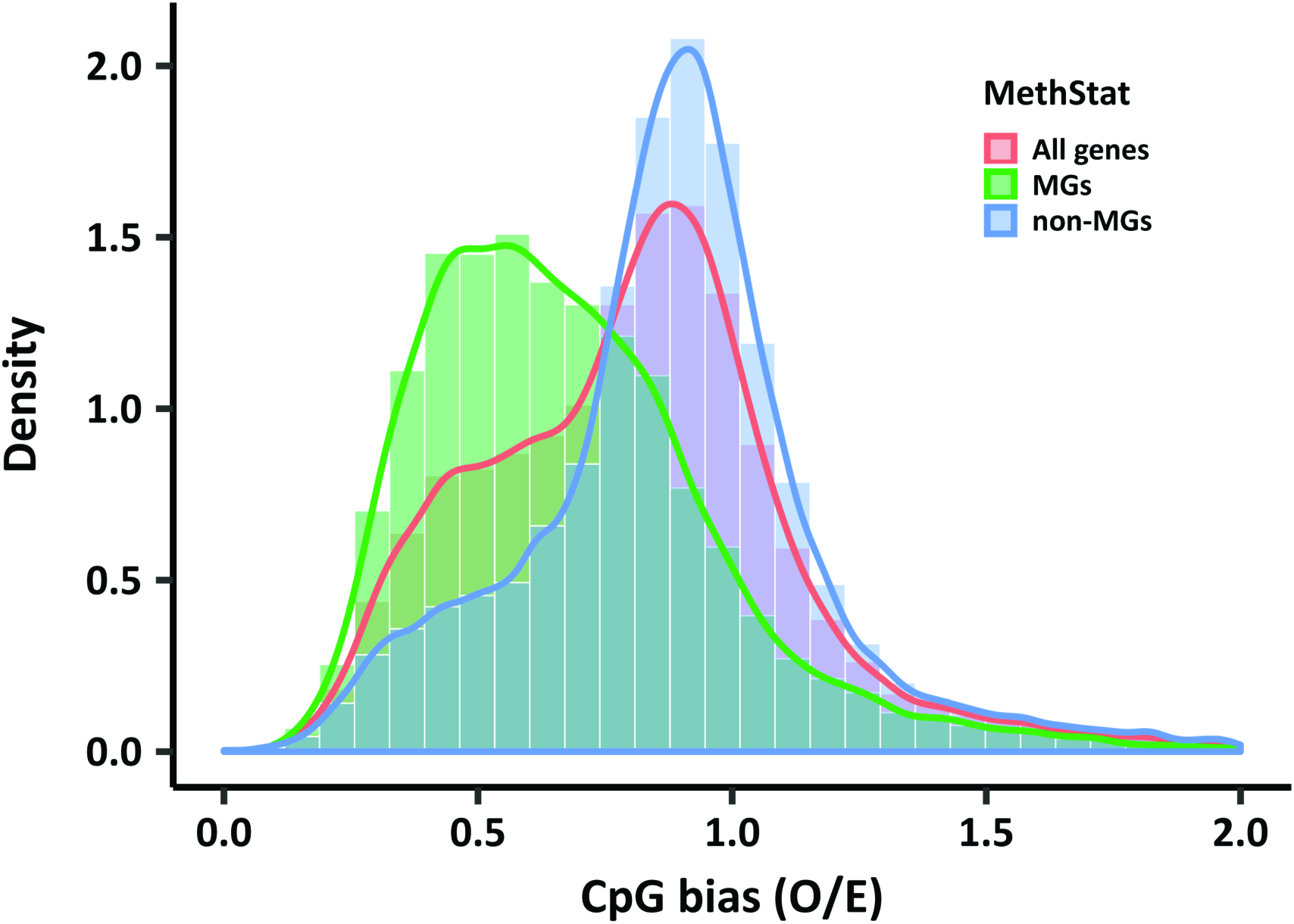
Methylated genes in Aiptasia have lower CpG O/E. CpG distribution of methylated genes (represented by red curve) peaks at around 0.5, which is lower than in unmethylated genes (represented by green curve) peaking at around 0.9. mC to T conversion skews the CpG O/E distribution of all genes as expected (represented by blue curve), but methylated and unmethylated genes still show a large overlap of their CpG O/E distributions. These results indicate that gene body methylation cannot be accurately inferred from CpG O/E in Aiptasia.

**Fig. S3.**
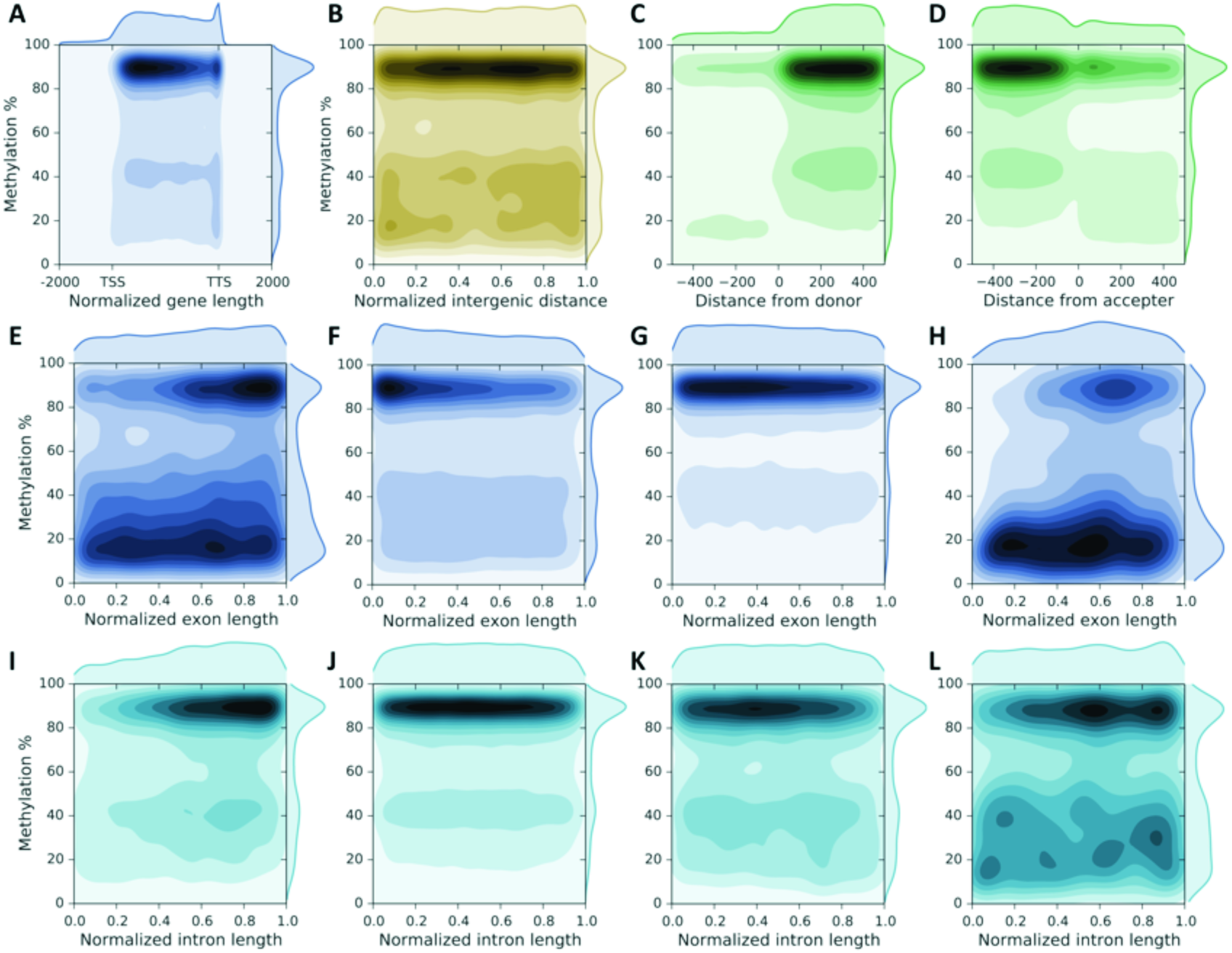
Methylation patterns. (**A**) DNA methylation is mainly located in the proximal part of gene bodies with slightly decreasing levels towards the end. (**B**) Methylation pattern over intergenic regions. (**C**) Methylation pattern around splice donor sites show increasing levels immediately after donor sites. (**D**) Methylation pattern around acceptor sites show decreasing levels immediately after splice acceptor sites. (**E**) Methylation pattern over initial exons show increasing methylation levels (3,147 exons with 35,885 methylation sites were used). (**F**) Methylation pattern over internal exons show decreasing methylation levels (7,977 exons with 139,009 methylation sites were used). (**G**) Methylation pattern over terminal exons show decreasing methylation levels (7,905 exons with 102,162 methylation sites were used). (**H**) Methylation pattern over introns from single-exon genes follow a similar trend as observed for multi exon genes with increasing methylation levels in the proximal and decreasing levels in the posterior part of the exon (298 exons with 4,735 methylation sites were used). (**I**) Methylation pattern over initial introns show increasing methylation levels (3,381 introns with 39,262 methylation sites were used). (**J**) Methylation pattern over internal introns maintain stable methylation levels (7,371 introns with 211,950 methylation sites were used). (**K**) Methylation levels over terminal introns decrease slightly (3,959 introns with 34,246 methylation sites were used). (**L**) Methylation levels over introns from one-intron genes change gently with initial increase followed by a decrease (1,055 introns with 10,709 methylation sites).

**Fig. S4.**
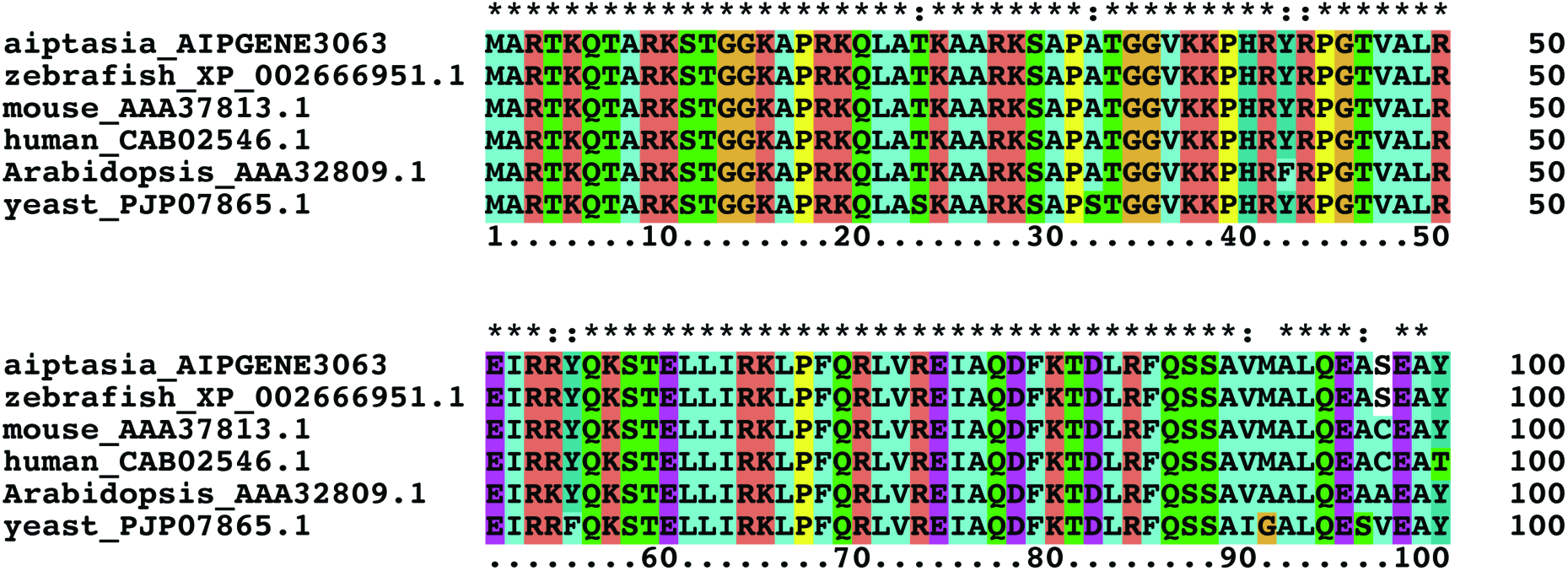
Sequence conservation of histone H3 homologs. Sequence conservation of Aiptasia histone H3 protein and histone H3 homologs from species for which antibody (ab9050, Abcam) has previously been validated. The N-terminal tail of Aiptasia H3 is identical to the fragment from the zebrafish *Danio rerio* (the first 100 amino acid fragment from human was used to produce this antibody).

**Fig. S5.**
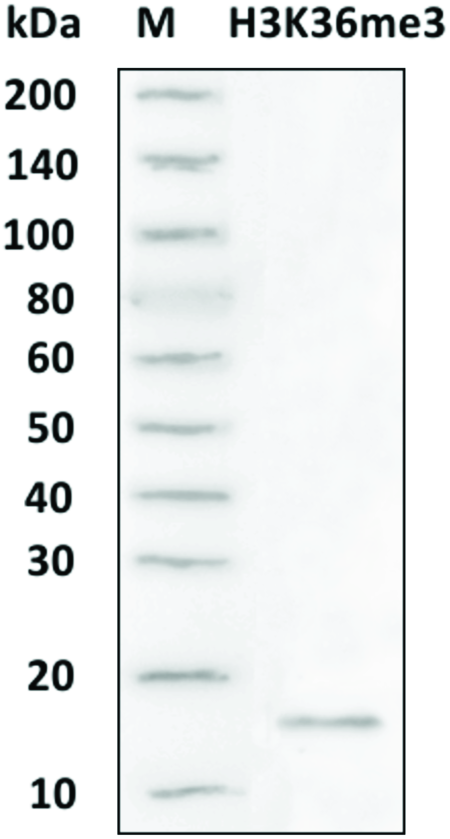
Western blot. Western blot result for antibody affinity validation, target band is 15kDa in size as expected from molecular weight analysis.

**Fig. S6.**
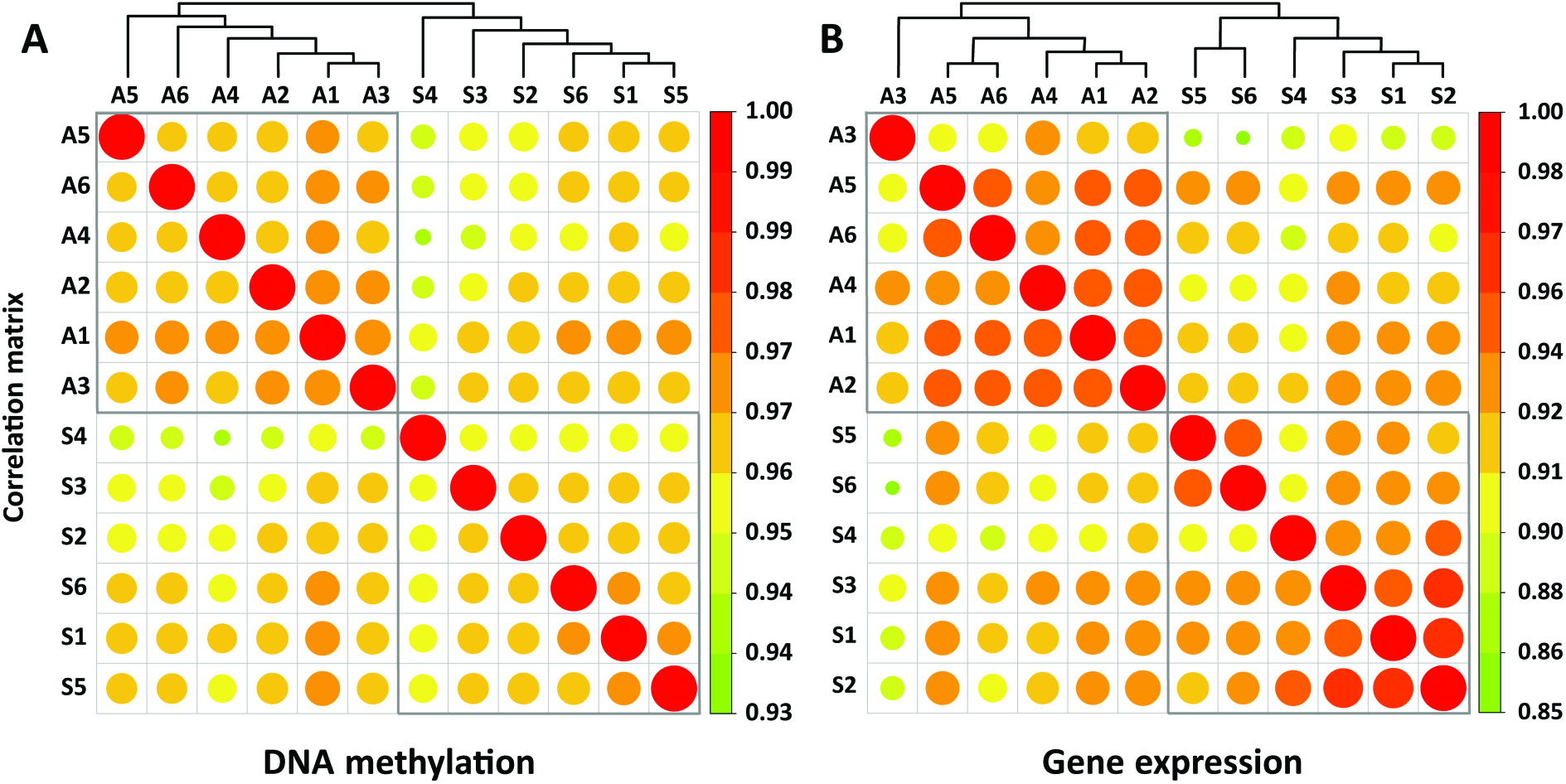
Correlation matrices of replicates. Correlation matrices of replicates based on median DNA methylation level of genes (**A**) and log gene expression values (base 2) (**B**). Replicates from the same treatments showed higher correlation and clustered together both based on DNA methylation as well as gene expression profiles, further supporting the findings obtained from the PCA analyses (figure 4) that changes in DNA methylation and expression are treatment specific.

**Fig. S7.**
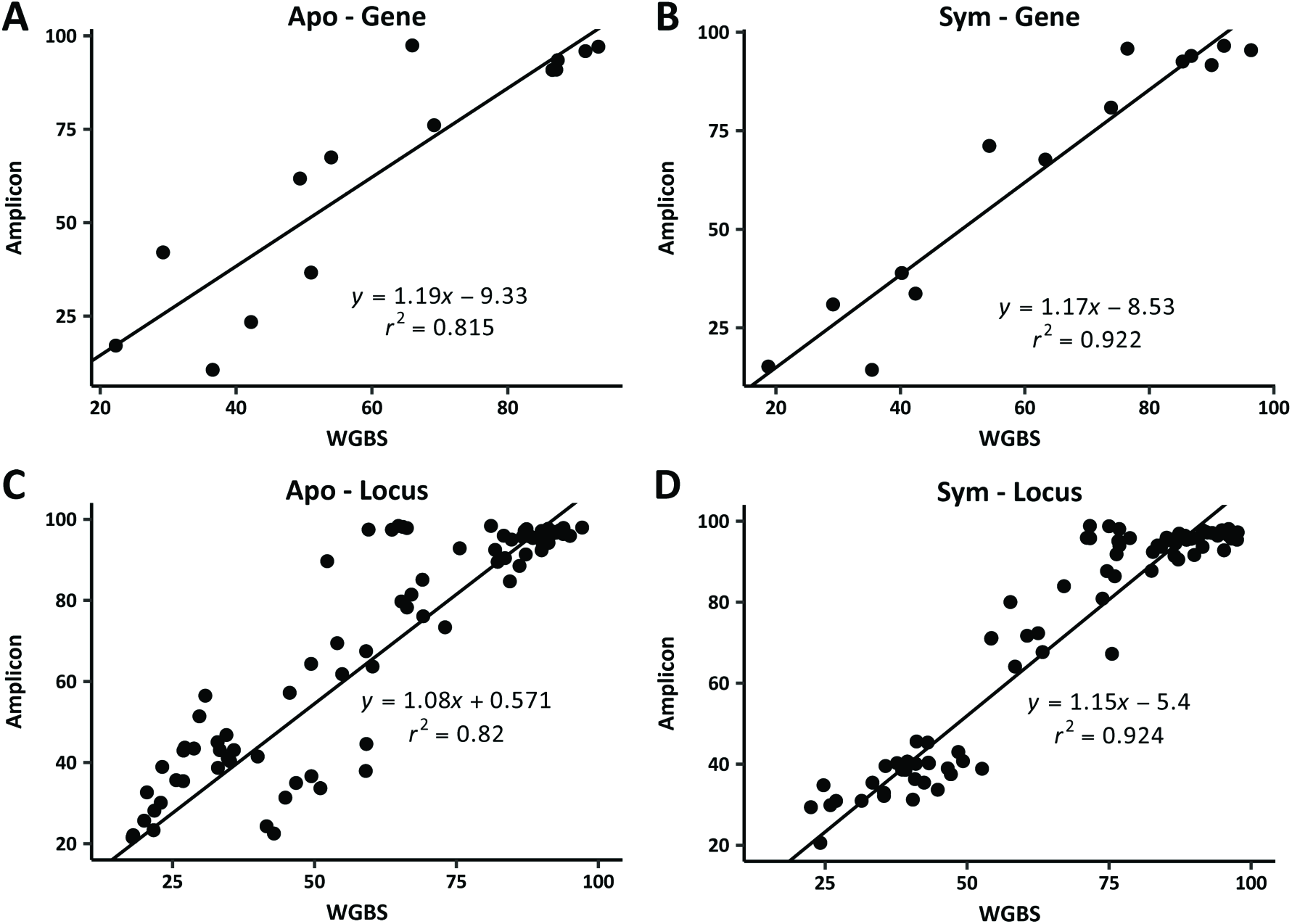
Validation of methylation level. Validation of methylation level using bisulfite PCR on selected genes. (**A**, **B**) validation of methylation level on genes (median methylation levels of methylated CpGs were used to represent genes). (**C**, **D**) validation of methylation level on locus (methylated CpGs). WGBS: whole genome bisulfite sequencing; Amplicon: MiSeq sequencing results of bisulfite PCR amplicons on selected genes.

**Fig. S8.**
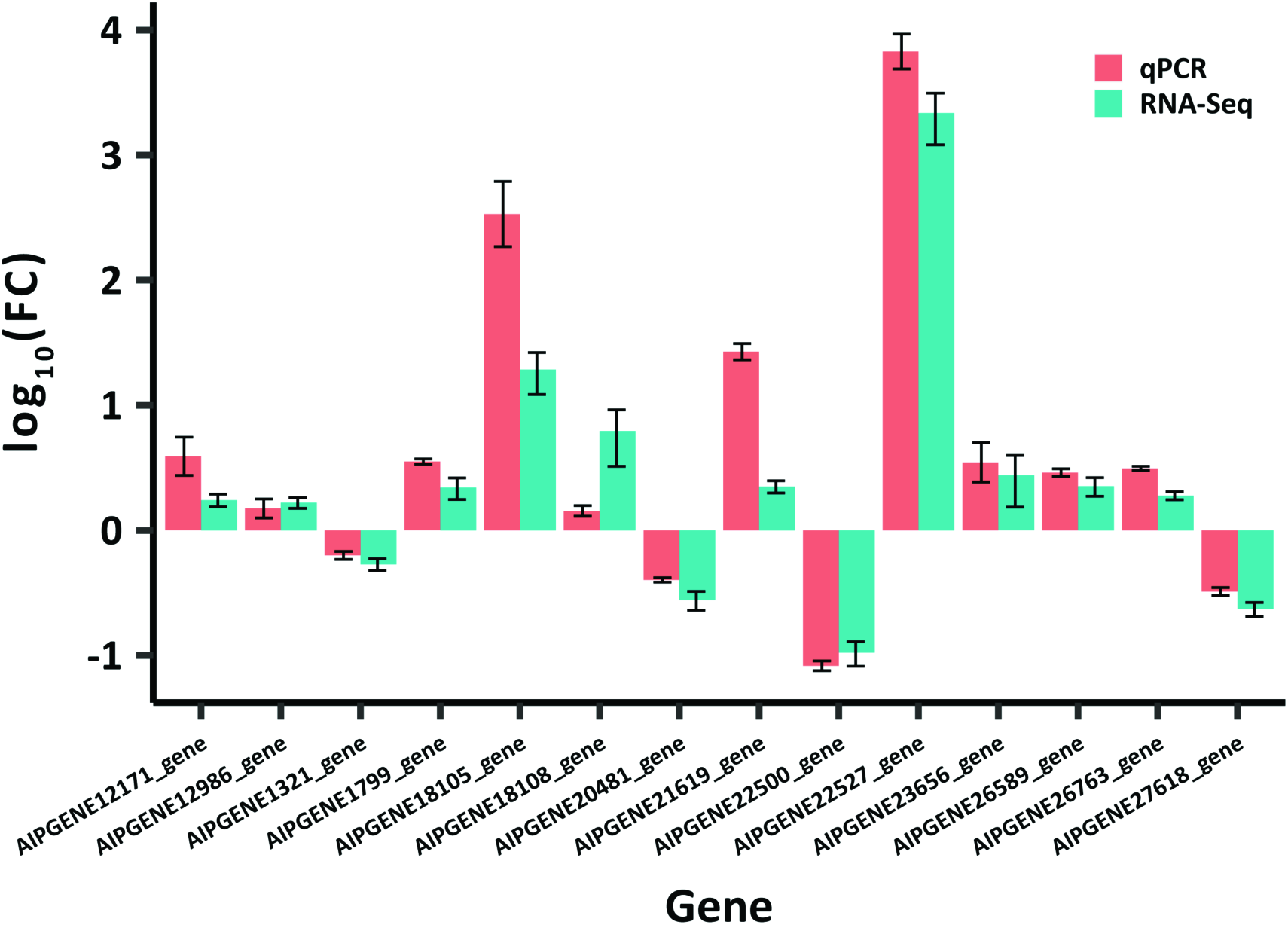
qPCR validation of gene expression levels. Validation of gene expression changes using qPCR. Expression levels are shown as log_10_(fold change). All genes show similar expression changes as determined by RNA-seq and q-PCR.

**Fig. S9.**
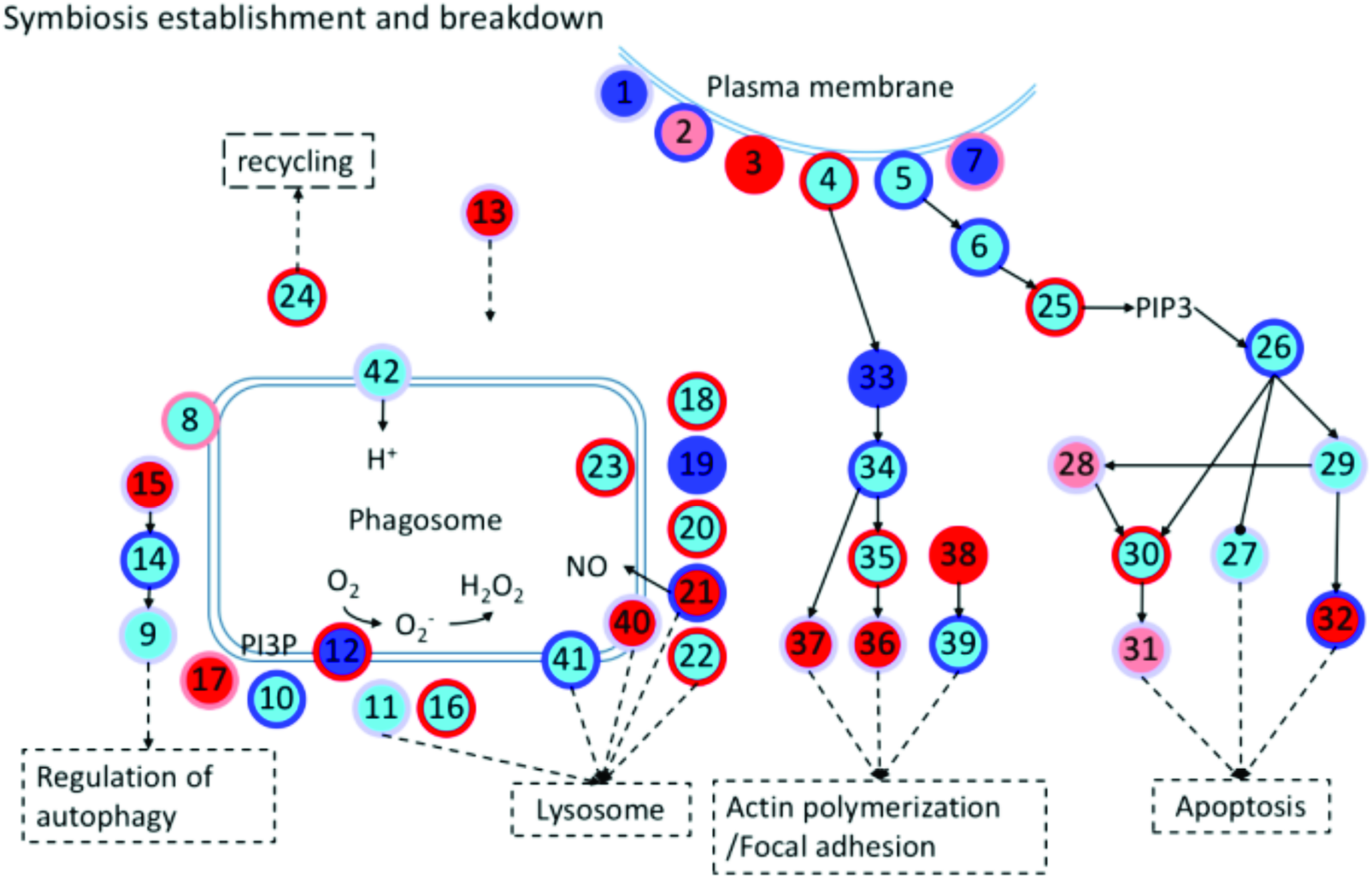
Schematic diagram of symbiosis establishment and breakdown associated genes. Every node represents a category of genes, and generally has multiple corresponding genes. The inside colors of nodes represent the expression changes of corresponding genes, including non-DEGs (cyan), up-regulated (red), down-regulated (blue) and up-and down-regulated DEGs (light red). The colors of node edges represent the methylation level changes of corresponding genes, including non-DMGs (light blue), hypermethylated (red), hypomethylated (blue) and hyper-and hypo-methylated DMGs (light red). Numbers in circles denote genes/proteins as detailed below.

1. Complement receptor

2. Scavenger receptor

3. C-type lectin

4. Integrin

5. Toll-like receptor

6. Ras-related C3 botulinum toxin substrate 1-rho family (RAC1)

7. Collagen

8. Vesicle-associated membrane protein (VAMP)

9. Autophagy-related protein 16 (ATG16)

10. Ras-related protein 5 (Rab5)

11. Ras-related protein 7 (Rab7)

12. NADPH oxidase (NOX)

13. Syntaxin 12

14. Autophagy-related protein 5 (ATG5)

15. Autophagy-related protein 10 (ATG10)

16. Programmed cell death 6-interacting protein

17. Sorting nexin (SNX)

18. Cytoplasmic dynein

19. Tubulin alpha chain (TUBA)

20. Tubulin beta chain (TUBB)

21. Nitric oxide synthase (NOS)

22. Lysosome-associated membrane glycoprotein/Cluster of differentiation (LAMP/CD)

23. Cathepsin L

24. Kinesin

25. Phosphatidylinositol 4,5-bisphosphate 3-kinase (PI3K)

26. RAC serine/threonine-protein kinase (AKT)

27. Bcl-2-antagonist of cell death (BAD)

28. TNF receptor-associated factor (TRAF)

29. Nuclear factor of kappa light polypeptide gene (NFKB)

30. Caspase 8 (CASP8)

31. Caspase 7 (CASP7)

32. Apoptosis regulator/Bcl-2 (BCL2)

33. Ras homolog (RHO)

34. Rho-associated protein kinase (ROCK)

35. Phosphatidylinositol 4-phosphate 5-kinase/Phosphatidylinositol 5-phosphate 4-kinase/Phosphatidylinositol 3-phosphate 5-kinase (PI4P5K/ PI5P4K/ PI3P5K)

36. Vinculin

37. Radixin

38. Profilin

39. Actin

40. CD63

41. Lysosomal-associated transmembrane protein

42. V-type proton ATPase

PI3P: phosphatidylinositol-3-phosphate

PIP3: Phosphatidylinositol (3,4,5)-trisphosphate

**Fig. S10.**
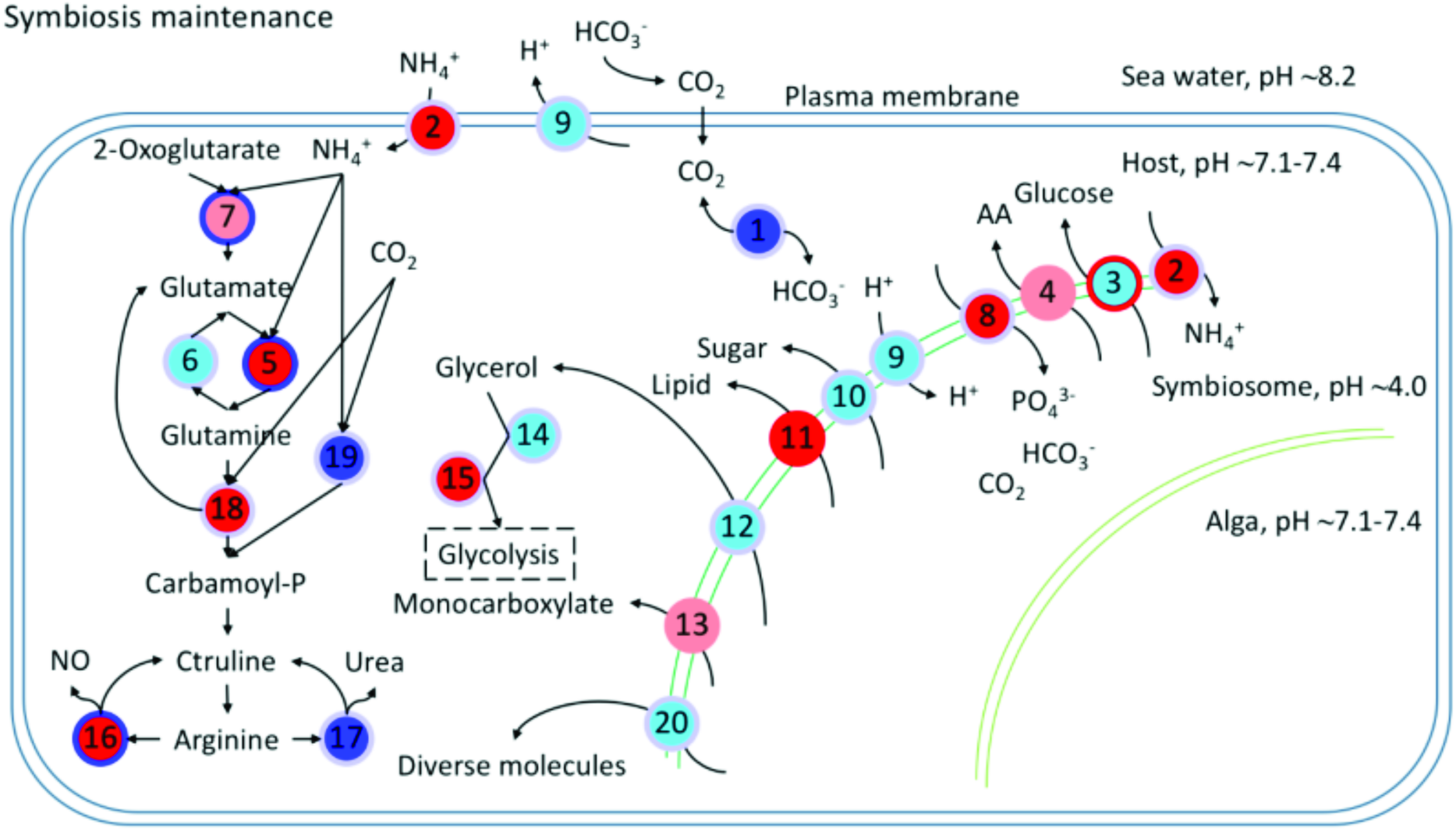
Schematic diagram of symbiosis maintenance associated genes. Every node represents a category of genes, and generally has multiple corresponding genes. The inside colors of nodes represent the expression changes of corresponding genes, including non-DEGs (cyan), up-regulated (red), down-regulated (blue) and up-and down-regulated DEGs (light red). The colors of node edges represent the methylation level changes of corresponding genes, including non-DMGs (light blue), hypermethylated (red), hypomethylated (blue) and hyper-and hypo-methylated DMGs (light red).

1. Carbonic anhydrase (CA)

2. Ammonium transporter

3. Glucose transporter

4. Amino acid transporter

5. Glutamine synthetase (GS)

6. Glutamate synthase

7. Glutamate dehydrogenase (GDH)

8. Phosphate transporter

9. V-type proton ATPase

10. Sugar transporter

11. Lipid transfer protein

12. Aquaporin 3 (Glycerol transporter)

13. Monocarboxylate transporter

14. Alcohol dehydrogenase

15. Aldehyde dehydrogenase

16. Nitric oxide synthase (NOS)

17. Arginase

18. Carbamoyl-phosphate synthase / Aspartate carbamoyltransferase / Dihydroorotase (CAD)

19. Carbamoyl-phosphate synthase (ammonia) (CPS1)

20. ABC transporter

